# Filling gaps on the diversity and biogeography of Chilean millipedes (Myriapoda: Diplopoda)

**DOI:** 10.1101/2022.06.11.495770

**Authors:** Antonio Parra-Gómez, Leonardo D. Fernández

**Author notes:** Corresponding author: Leonardo D. Fernández.

## Abstract

Research on the diversity and biogeography of Chilean millipedes (Diplopoda) represents a severe gap in knowledge. To reduce this gap we conducted a study to: (1) investigate the state of knowledge of millipede diversity, and (2) assess the pattern and causes underlying the latitudinal diversity gradient in Chilean millipedes. After combining the number of described species with those that have not yet been formally described, we concluded that there are 95 native millipede species in Chile. A diversity estimate suggested that in the future this number could increase to 125 or 197 species. However, this estimate is based on limited data. Therefore, the number of millipede species inhabiting Chile probably exceeds our estimate. Consistently, rarefaction-extrapolation curves revealed that we have not yet recorded a substantial fraction of millipede diversity and that increased sampling effort will reveal the presence of a greater number of millipede species in Chile. Most millipede species exhibited narrow geographic ranges in Chile. The north-south distribution of their species richness followed a bell-shaped latitudinal gradient of diversity, i.e. diversity peaked at the temperate climate of central Chile and then decreases towards the arid and polar climates of northern and southern Chile, respectively. The causes underlying this biogeographical pattern were water availability, ambient energy input and climate stability. This finding provided support for two of the five biogeographic hypotheses we tested: water-energy balance and climate stability. Thus, Chilean millipedes were more diverse at sites that exhibit warm and humid (temperate) climates for much of the year.

## 1. Introduction

Diplopoda is the third largest class of terrestrial arthropods after Insecta and Arachnida (Golovatch and Kime 2009). Its diversity is close to 12,000 species although it is estimated that there could be as many as 80,000 species worldwide (Golovatch and Kime 2009). Although diplopods are commonly referred to as millipedes, their representatives have only hundreds (or fewer) pairs of legs distributed along their body segments (Adis 2002). So far the only known exception to this rule is the species *Eumillipes persephone*, which boasts 1,306 pairs of legs (Marek et al. 2021). Found on all continents except Antarctica, millipedes inhabit a wide range of terrestrial biomes although they are most diverse in those with warm and humid climates, such as temperate forests (Minelli 2015).

The diversity and distribution of millipedes is poorly known compared to that of other historically better studied metazoans (Sierwald and Bond 2007). These gaps in knowledge are severe in Chile (Golovatch 2014; Parra-Gómez submitted), a long and narrow country in southwestern South America (Fig. 1a). Chile stretches from 17° S to 56° S covering c. 4,300 km from north to south and yet it only averages 177 km from east to west. Chile’s extensive latitudinal distribution is coupled with a strong north–south climatic gradient. Broadly speaking, the north at low latitudes is arid, the centre at mid-latitudes is temperate, while the south at high latitudes is polar (Beck et al. 2018).

**Figure 1.**
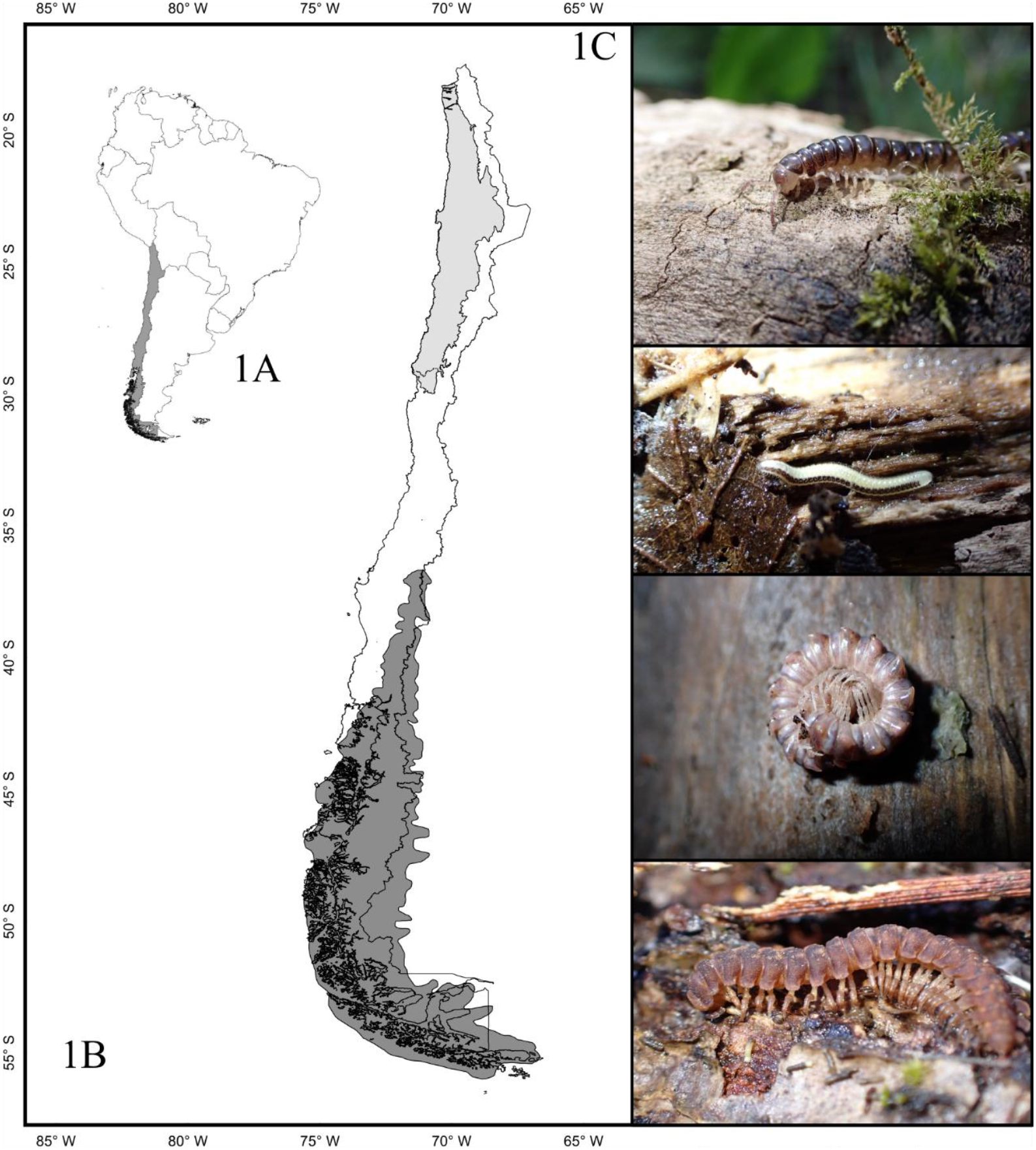
Map and millipedes of Chile. (A) Chile in southwestern South America stretches from ∼17°S to 56°S. (B) During the Pleistocene, a series of geological events (e.g. the uplift of the Andes) and climatic changes (e.g. the Last Glacial Maximum) shaped the Atacama Desert in the north (area shaded in light grey) and the Patagonian ice sheet in the south (area shaded in dark grey) in Chile. Since then, the Atacama Desert has favoured an arid climate at low latitudes while at the end of the Last Glacial Maximum the retreat of the Patagonian Ice Sheet set up a polar climate at high latitudes. The mid-latitudes of central Chile exhibit a temperate climate and acted as a refuge for biota during the occurrence of the historical contingencies described. **c**. Examples of millipede species endemic to Chile. From top to bottom: *Tsagonus* aff. *valdiviae* Chamberlin, 1957; *Siphonotus parguaensis* Mauriès and Silva, 1971; *Monenchodesmus inermis* Silvestri, 1903 and *Mikroporus granulatus* Attems, 1898.

The Chilean north–south climatic gradient emerged during the Pleistocene as a consequence of desertification and glaciation events that modified the original warm and humid conditions of northern and southern Chile respectively (Fig. 1b) (Gregory-Wodzicki 2000; Davies et al. 2020). Driven by these historical contingencies many northern and southern mesophile taxa became extinct or migrated to the temperate climate of central Chile, where they subsequently diversified (Villagrán and Hinojosa 1997). Today, the diversity of most taxa peaks in mid-latitudes and then decreases towards low and high latitudes (Villagrán and Hinojosa 1997; Samaniego and Marquet 2009; Fernández et al. 2015; Moreno et al. 2021; Campello-Nunes et al. 2022).

Chile is often referred to as a biogeographic island because its political boundaries overlap with natural barriers to species dispersal. To the north Chile is bordered by the Atacama Desert, to the east by the Andes Mountains, to the west by the Pacific Ocean and to the south by the end of the South American continent (Figs 1a-b). The Chilean biota therefore evolved practically in isolation and today Chile is a well-known hotspot of biodiversity and endemism for macro- and microscopic organisms (Arroyo et al. 2004; Fernández et al. 2015; Campello-Nunes et al. 2022), including millipedes (Parra-Gómez submitted).

The insular condition of Chile and the historical contingencies experienced by its territory probably favoured the occurrence of a particular millipede diversity (Fig. 1c). Chilean millipedes are mainly Gondwanan relicts (Shelley and Golovatch 2011). They share more taxonomic and possibly evolutionary affinities with millipedes from South Africa, Australia, Tasmania, New Caledonia and New Zealand than with other South American taxa (Golovatch 2014). Families such as Dalodesmidae and Eudigonidae are widely represented in Chile and only marginally in Argentina, while the family Iulomorphidae is only found in Chile.

The goal of the present study is to investigate the state of knowledge and biogeography of Chilean millipedes. Specifically, we investigated the temporal variation in the number of published studies, the number of sites investigated, and the number of new species reported in Chile over the last 175 years. We also constructed rarefaction, extrapolation and asymptotic diversity curves to assess the number of species that have not yet been described in Chile. Finally, we tested six biogeographic hypotheses (Table 1) to investigate the pattern and causes underlying the latitudinal diversity gradient in Chilean millipedes.

**Table 1.** Ecological hypotheses tested with respect to the diversity of native millipedes in Chile and their respective predictions and predictors.

| Hypothesis | Prediction | Predictors | Reference examples |
| --- | --- | --- | --- |
| 1 Species-energy | Diversity increases with energy inputs (including plant productivity) | UVB, NPP, NDVI, PET, AMT, MTW, MTQ, MWQ, MCQ | Currie (1991), Evans et al. (2005) |
| 2 Water availability | Diversity increases with water inputs | APR, PDM, PWQ | Rodríguez et al. (2005), Samaniego and Marquet (2009) |
| 3 Water energy-balance | Diversity increases with energy and water inputs | AET, UVB, NPP, NDVI, PET, APR, PDM, PWQ | Hawkins et al. (2003), Rodríguez et al. (2005), Fernández et al. (2016) |
| 5 Climatic variability | Diversity increases in climatically stable areas | ISO, TS, MDR, TAR, MTC, | Currie (1991), Kerr and Packer (1997) |
| 6 Habitat heterogeneity | Diversity increases with habitat heterogeneity | ELE | Pianka (1966), Kerr and Packer (1997) |

UVB: Ultraviolet Radiation B, NPP: Net Primary Productivity, NDVI: Normalized Difference Vegetation, PET: Potential Evapotranspiration, AMT: Annual Mean Temperature, MTW: Max Temperature of Warmest Month, MTQ: Mean Temperature Of Wettest Quarter, MWQ: Mean Temperature of Warmest Quarter, MCQ: Mean Temperature of Coldest Quarter, APR: Annual Precipitation, PDM: Precipitation of Driest Month, PWQ: Precipitation of Warmest Quarter, ISO: Isothermality, TES: Temperature Seasonality, MDR: Mean Diurnal Range, TAR: Temperature Annual Range, MTC: Min Temperature of Coldest Month, ELE: Topographic Elevation

## 2. Methods

### 2.1. Data source

The database used in this study consists of 95 native species recorded in continental Chile between 1847 and 2022. We constructed the database based on species records obtained by the first author and species records included in a recent checklist of Chilean millipedes (Parra-Gómez submitted). Our database is the most up-to-date and comprehensive database for millipedes in Chile. We excluded from the original database (109 species) a total of eight exotic species and six *nomina dubium* species. We used our database to construct a binary matrix (i.e. columns as samples and rows as taxa) on which all subsequent statistical analyses were based.

### 2.2. State of knowledge of Chilean millipedes

We used three complementary approaches to investigate the state of knowledge of Chilean millipedes following Fernández et al. (2015) and Campello et al. (2022).

First, we estimated the number of published studies, the number of new sites sampled, the number of new species reported, and the cumulative number of new species reported in each decade from the 1840s onwards. We then correlated each indicator against time to investigate its trend over years using the R package spdep version 1.1–12 (Bivand et al. 2013).

Second, we estimated a rarefaction curve to investigate whether the sampling effort invested in Chile (measured as the number of sites investigated between the 1840s and 2020s) has contributed to the completion of the checklist of Chilean millipede species. If the rarefaction curve reaches a plateau, we will conclude that the investment of additional sampling effort will not reveal a substantial number of new species in Chile. If the rarefaction curve does not reach a plateau, then we will conclude that the investment of additional sampling effort will reveal new species in Chile. We also estimated an extrapolation curve to investigate whether an increase in sampling effort will contribute to the completeness of the checklist of Chilean millipede species. If the rarefaction curve reaches a plateau, we will conclude that an increase in sampling effort will contribute to completing the checklist of Chilean millipede species. If the rarefaction curve does not reach a plateau, we will conclude that a significant increase in sampling effort is needed to complete the checklist of Chilean millipede species. We estimated the rarefaction and extrapolation curves based on the approach proposed by Chao et al. (2014). The rarefaction curve was estimated using the number of sites surveyed in Chile from the 1880s onwards as a proxy for sampling effort (n = 140 sampling sites). The extrapolation curve was estimated by doubling the sampling effort (n = 280 sampling sites). The rarefaction and extrapolation curves and their lower and upper confidence limits (95% CI) were estimated based on 10,000 bootstrap replicates in the R package iNEXT version 2.0.20 (Hsieh et al. 2016).

Third, we constructed an asymptotic diversity profile to estimate the number of species we have yet to discover at the sites we have so far surveyed in Chile. We estimated the asymptotic diversity profile and its upper and lower 95% confidence intervals (10,000 bootstrap replicates) based on the method proposed by Chao and Jost (2015) implemented in the R package iNEXT version 2.0.20 (Hsieh et al. 2016).

### 2.3. Millipede biogeography

We used a range interpolation approach to standardise species richness and reduce the effects of spatial sampling biases (McCain 2009; Fernández et al. 2022). Range interpolation assumes that species have continuous geographic ranges between their lowest and highest latitudinal occurrences. Standardised species richness was subsequently used to investigate the latitudinal diversity gradient, the size of geographic ranges and beta diversity of Chilean millipedes.

To investigate the latitudinal diversity gradient we divided Chile into bins of 3° latitudinal bands. We then counted the number of species recorded in each latitudinal bin and correlated species richness against latitude by fitting linear and non-linear functions in PAST version 4.09 (Hammer et al. 2001). We used the Akaike information criterion to select the model that best fitted the data.

To investigate the distribution ranges of millipede species we plotted and classified the latitudinal distribution of each species into one of the following categories: (a) species with small distributions, occurring only within two latitudinal bands; (b) species with narrow-medium ranges of distribution, ranging from three to 10 latitudinal bands; (c) species with medium-large distributions, ranging from 11 to 19 latitudinal bands; and (d) species with large distributions, ranging from 20 to 39 latitudinal bands.

We investigated the relationship between species richness and 25 environmental variables frequently used as proxies for ecological hypotheses proposed to explain the occurrence of latitudinal diversity gradients (Table 1). We obtained the environmental variables from various sources such as WorldClim, among others. Based on scatter plots for all variable pairs (Draftsman Plot) we log (x + 1) transformed all variables and removed highly correlated variables to avoid skewed trends. This process resulted in 19 environmental variables (Table 1). We then normalized all selected variables to compare variables with different unit measures (Clarke et al. 2005). To investigate the relationship between these 19 variables and millipede richness we conducted a BioEnv procedure, which used a multiple regression approach to determine which environmental variables best explain the latitudinal diversity gradient in millipedes (Clarke and Ainsworth 1993). We estimated BioEnv and its statistical significance (1,000 permutation) in PRIMER version 6 (Clarke and Gorley 2006).

To investigate beta diversity or the latitudinal variation in species composition we estimated beta diversity (β_SOR_) as well as its underlying additive components, i.e. spatial turnover (β_SIM_) and nestedness (β_SNE_), as described by Baselga (2012). In brief β_SOR_ is a measure of (di)similarity based on Sørensen’s index and represents a global metric of beta diversity. β_SIM_ is a measure of (di)similarity based on Simpson’s index and describes the fraction of beta diversity that corresponds only to the spatial turnover of species. β_SNE_ is estimated as the difference between β_SOR_ and β_SIM_ and describes the spatial loss of diversity among sites. These metrics range from zero (perfect similarity) to one (perfect dissimilarity). We estimated β_SOR_, β_SIM_ and β_SNE_ using the R package betapart (Baselga and Orme 2012).

We also conducted an analysis of similarity (ANOSIM) to investigate the variation in species composition between arid, temperate and polar climates at low, mid and high latitudes, respectively. ANOSIM compares the mean of ranked dissimilarities between groups to the mean of ranked dissimilarities within groups based on the R–statistic. An R–statistic close to “1” suggests dissimilarity between groups, an R–statistic close to “0” suggests an even distribution of high and low ranks within and between groups, while an R–statistic below “0” suggest that dissimilarities are greater within groups than between groups. The ANOSIM and significance value of the R–statistic were estimated based on 1000 permutations in PRIMER version 6 (Clarke and Gorley 2006).

## 3. Results

### 3.1. State of knowledge of Chilean millipedes

We recorded a positive and significant correlation between time and the number of published studies, the number of sites sampled, the number of new species reported, and the cumulative number of new species reported in each decade (Fig. 2). These results suggest that the knowledge on Chilean millipedes has increased (albeit modestly) from the 1840s onwards.

**Figure 2.**
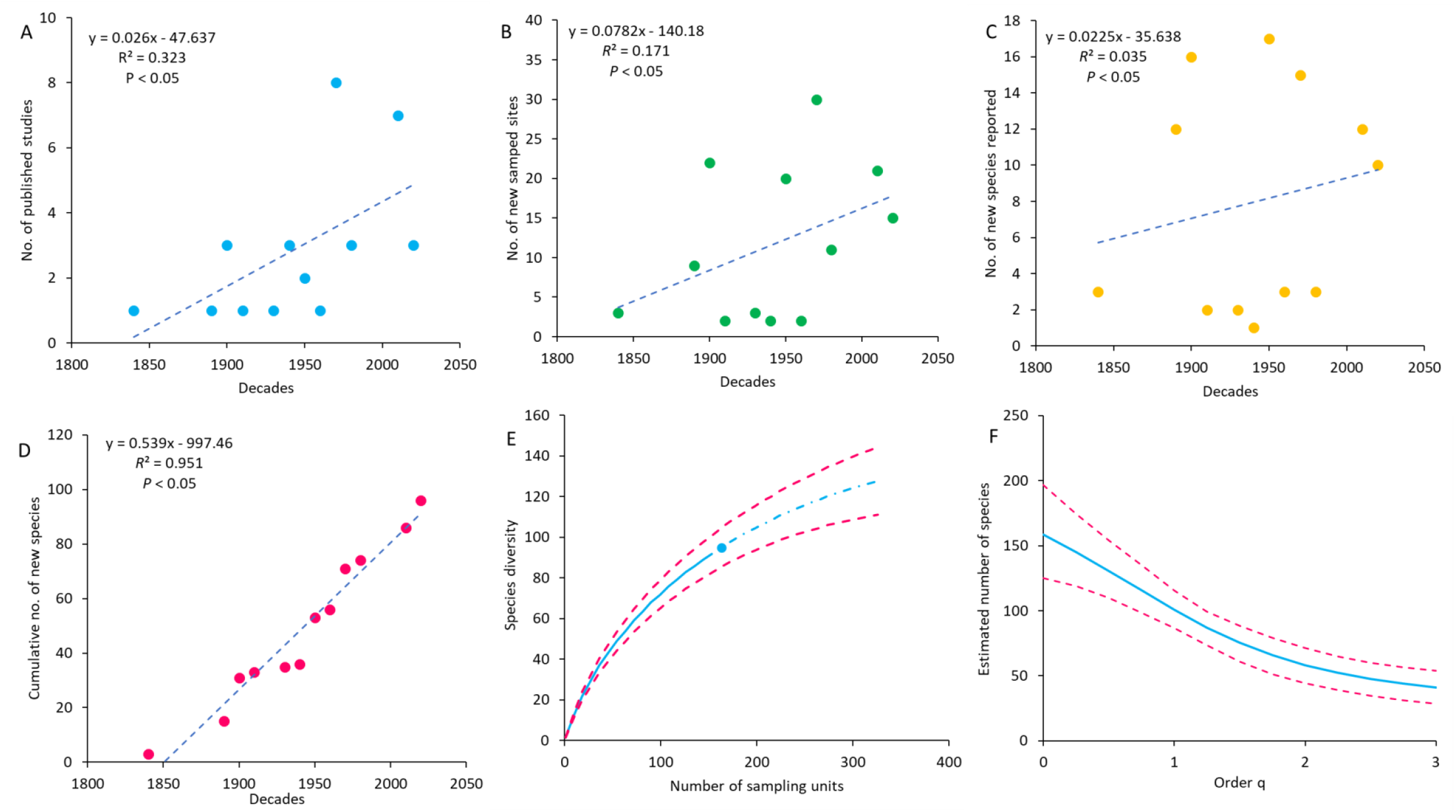
State of knowledge of Chilean millipedes. (A) number of studies published per decade. (B) number of new sites sampled per decade. (C) number of new species reported per decade. (D) cumulative number of new species per decade. (E) rarefaction curves (solid trend line) and extrapolation curves (dash-dotted line). The point between both curves represents the number of sites where millipedes have been sampled in Chile. (F) Asymptotic estimate of millipede diversity in Chile (solid trend line).

Our analysis revealed that between 1847 and 2022 (175 years) native millipede species were described or reported in 30 studies. The number of published studies exhibits two peaks over time, i.e. one in the 1970s and another in the 2010s with eight and seven published studies, respectively. The remaining decades exhibit between one and three published studies. While we recorded an increase in the number of studies on Chilean millipedes over time, we also noted extended periods without published studies. These include the periods between 1848 and 1897 (49 years); 1906 and 1915 (nine years) and more recently between the 1989 and 2011 (22 years) (Fig. 2a).

The review of the 30 published studies on Chilean millipedes revealed that researchers have explored the diversity of these invertebrates at 140 unique sites. Eleven studies (37%) are based on a single study site, four (13%) on three sites, three (10%) on two sites, three (10%) on eight sites, two (7%) on 13 sites, two (7%) on six sites, two (7%) on five sites, one (3%) on 19 sites, one (3%) on 11 sites and one (3%) on nine sites. The number of new sites sampled exhibits a peak in the 1970s with 30 new sites sampled. The 1900s, 1950s and 2010s also stand out with 22, 20 and 21 new sites sampled. The number of new sites sampled during the remaining decades range between 1 and 11 sites. Overall our analysis suggests that, even though there are extended periods without published studies, the number of new sites investigated has increased over time, particularly after the 1900s (Fig. 2b).

The number of new millipede species also increases significantly over time although the correlation coefficient is low. This is because the number of new species reported varies significantly from decade to decade, with values ranging from one to 17 species. The highest number of new species was recorded in the 1950s (17 species), while the lowest number of species was recorded in the 1910s and 1940s (1 species each time) (Fig. 2c).

Although the number of new species reported varies significantly between decades, the cumulative number of new species has increased exponentially over time. Between the 1840s and 2020s the number of native species known for Chile has increased from three to 95 (including species that have yet to be formally described; Parra-Gómez, personal observation). This suggests that conducting further studies and exploring more sites will reveal new species in Chile (Fig. 2d).

Rarefaction and extrapolation curves confirmed the above result. The rarefaction curve did not reach a plateau suggesting that the sampling effort has been insufficient to record a substantial number of the species inhabiting Chile. So, future studies will add new species to the checklist of Chilean millipedes. The extrapolation curve also did not reach a plateau, suggesting that doubling the sampling effort would also not contribute to recording a significant fraction of the millipede species present in Chile. Therefore, we need to invest a greater sampling effort over time to complete the checklist of Chilean millipedes (Fig. 2e).

The diversity estimate suggests that together the sites surveyed between the 1840s and 2020s harbour at least 158 native species (considering 95% lower and upper confidence intervals of 125 and 197 species). Thus, our analysis suggests that we have missed at least one-third of the millipede species that actually inhabit the sites surveyed between the 1840s and 2020s. Therefore a significant increase in sampling effort could reveal up to 102 additional species (Fig. 2f).

### 3.2. Millipede biogeography

Millipedes exhibit a bell-shaped (unimodal) latitudinal diversity gradient in Chile. Thus, species richness exhibits a peak in the mid-latitudes (central Chile) and then decreases towards low (northern Chile) and high (southern Chile) latitudes (Fig. 3a).

**Figure 3.**
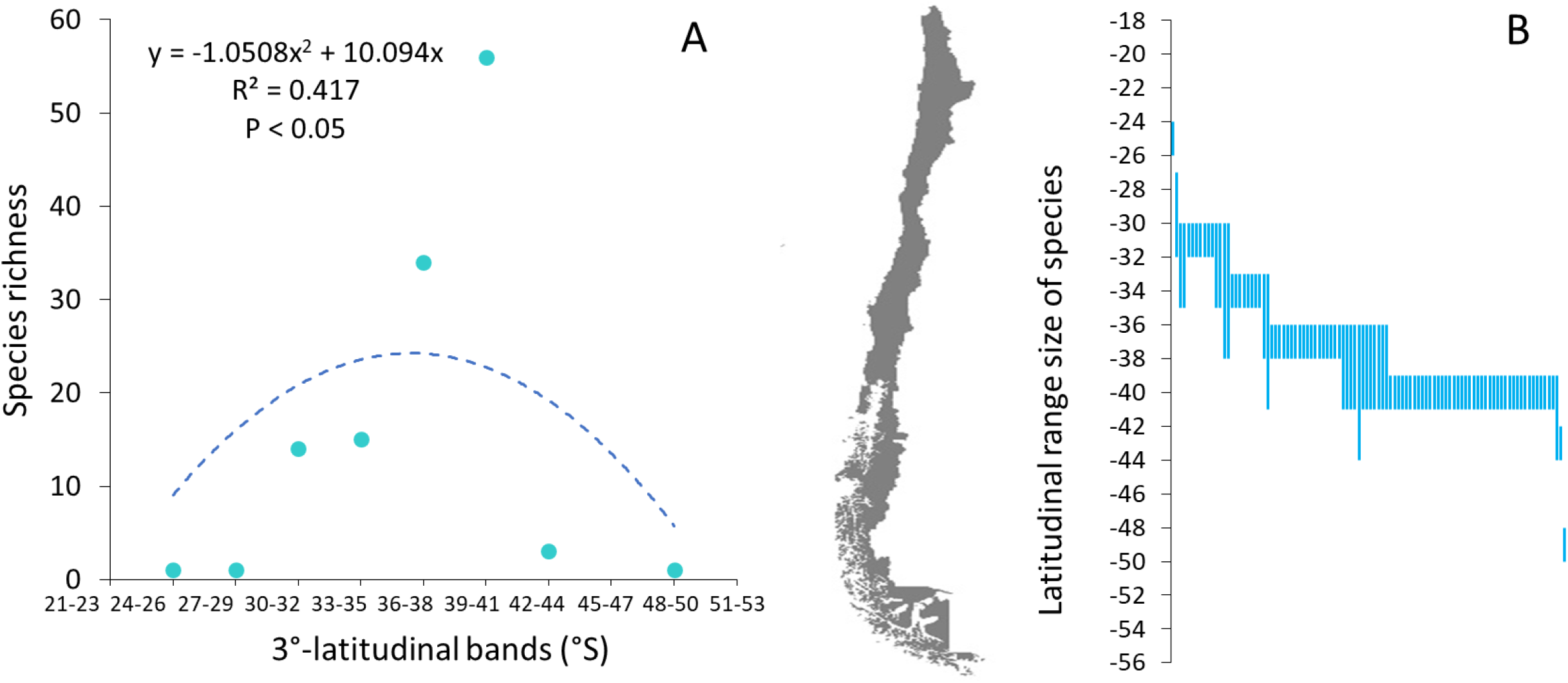
Biogeography of Chilean millipedes. (A) Bell-shaped latitudinal diversity gradient. The second-degree polynomial function (segmented line) reveals that millipede diversity peaks in central Chile and then decreases significantly towards northern and southern Chile. (B) Geographic ranges of Chilean millipedes. The geographic ranges of these invertebrates are narrow or narrow-medium and are mainly concentrated in central Chile.

Analysis of geographic range size revealed that most millipedes (78 species) have narrow geographic ranges, while the rest (28 species) have narrow-medium ranges (Fig. 3b). We did not record species with medium and medium-large geographic ranges in Chile. Our analysis also revealed that the diversity peak observed in mid-latitudes is a product of the accumulation (and overlap) of species with narrow geographic ranges in central Chile.

The bell-shaped latitudinal diversity gradient of Chilean millipedes is positively and significantly correlated with a subset of five environmental variables, including Ultraviolet Radiation B (UVB), Normalised Difference Vegetation (NDVI), Annual Precipitation (APR), Mean Diurnal Range (MDR) and Isothermality (ISO) (BioEnv, *R* = 0.787, *p* = 0.01). UVB and NDVI are proxies for energy, APR is a proxy for water availability, while MDR and ISO are proxies for climatic stability. The observed correlation between millipede diversity and these proxies lends support to the water–energy balance and the climate stability hypotheses. Thus, our results suggest that millipede diversity is high in mid-latitudes because they exhibit a continuous trade-off between water availability and ambient energy inputs throughout the year.

Millipedes have a high beta diversity in Chile (β_SOR_ = 0.94; Fig. 4). The most important underlying phenomenon was species turnover (β_SIM_ = 0.78; Fig. 4), suggesting high latitudinal variation in species composition. Nestedness was less important (β_SNE_ = 0.16; Fig. 4), suggesting that few species co-occur latitudinally in Chile. We also observed that species composition varies significantly between the arid, temperate and polar climates of low, mid and high latitudes, respectively (ANOSIM, Global *R* = 0.395, *p* = 0.043).

**Figure 4.**
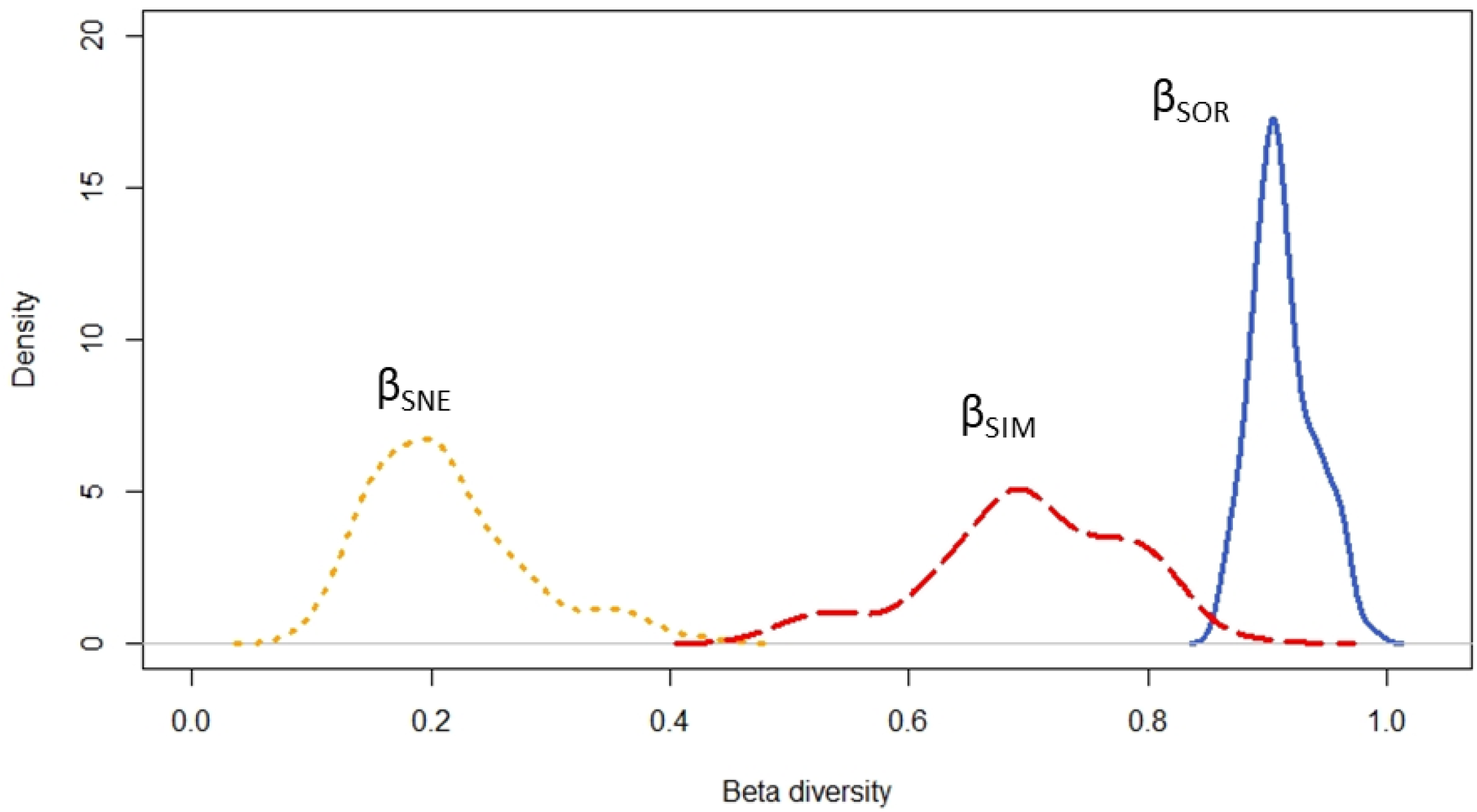
Beta diversity or latitudinal variation of Chilean millipede species composition. The curves in the kernel density plot show the distribution of observed values for beta diversity (β_SOR_, solid curve) and its two additive components, spatial turnover (β_SIM_, segmented curve) and nestedness (β_SNE_, dotted curve). According to this analysis beta diversity is mainly produced by millipede species turnover (β_SIM_) among northern (arid), central (temperate) and southern (polar) Chile.

## 4. Discussion

Millipedes are a poorly known group in Chile and represent a severe gap in knowledge for myriapodology worldwide (Golovatch 2014; Parra-Gómez submitted). In this study we investigated the state of knowledge, as well as the patterns and causes underlying the spatial distribution of their diversity across an extensive latitudinal gradient. To our knowledge, our study represents the first attempt to investigate the diversity and biogeography of these organisms in Chile.

### 4.1. State of knowledge of Chilean millipedes

Our analyses revealed that sampling effort and the number of published studies have increased, albeit only modestly, over the last 175 years. Both indicators are particularly low between the mid-19th and mid-20th centuries, probably because during that period it was very difficult to obtain samples from Chile. Researchers from Europe and North America travelled months to get to Chile and usually the samples did not arrive in good condition back to the laboratory (Certes 1889; Jung 1942). Other times researchers did not travel to Chile but analysed limited numbers of samples collected and granted by colleagues (Attems 1898; 1903) or scientific expeditions (Chamberlin, 1957). These limitations surely hindered the sampling effort, as well as the number and frequency of publications during that period. In fact, there is a gap of 51 years between the publication of the first study (Gervais 1847) and the second study (Attems 1898) on Chilean millipedes.

The sampling effort and the number of publications increased notably during the second half of the 20th century thanks to the contributions of the Chilean myriapodologist Francisco Silva. This researcher remained active for 13 years until his premature death in the 1990s (Silva et al. 1968; Demange and Silva 1971a, 1971b; Mauriés and Silva 1971; Silva and Vivar 1973, 1974; Silva and Sáiz 1975; Demange and Silva 1976a, 1976b; Urzua and Silva 1981). The studies of Krabbe (1982) and Shear (1988) are also added to this period. Finally, after a hiatus of 24 years, new studies on Chilean millipedes were published (Korsos and Read 2012; Golovatch 2014; Spelda 2015; Mesibov 2017; Short and Vahtera 2017; Vega-Román et al. 2019; Parra-Gómez and Faúndez 2021; Parra-Gómez 2022). Taken together, these studies represent a significant increase in sampling effort by exploring the diversity of Chilean myriapods at 36 new sites.

In Chile, there are 96 native millipede species. This value includes 68 described species in the last 175 years, species that have not yet been formally described and subspecies that will be promoted to species in the near future (Parra-Gómez submitted). However, the number of millipede species inhabiting Chile is possibly higher. Based on available data we estimated the diversity of native Chilean millipedes to be between 125 and 197 species. These values represent between two and three times the number of described species.

Although we used a robust method to assess millipede diversity (see Chao et al. 2015), we believe that we have underestimated the diversity of Chilean millipedes. Our estimate is based on diversity data collected at 140 sites, which represents a small fraction of the area included within Chile’s extensive latitudinal and altitudinal gradients. Each of these gradients contains thousands of sites where millipede diversity has never been investigated. In turn, each of these sites represents a myriad of microhabitats with local environmental conditions that could contain an imponderable number of native species unknown to Chile and science (Fernández 2011). Thus, the number of millipede species that have not yet been recorded in Chile probably exceeds our estimate. Our rarefaction and extrapolation curves suggest that we need to significantly increase sampling effort to record these species. Fortunately, there is renewed interest in investigating the diversity and distribution of Chilean millipedes (Vega-Román et al. 2019; Parra-Gómez and Faúndez 2021; Parra-Gómez submitted, present study). Therefore, our knowledge about Chilean millipedes could increase significantly in the coming years.

### 4.2. Millipede biogeography

After accounting for spatial bias in sampling effort we observed that millipede species richness follows a bell-shaped latitudinal diversity gradient, i.e. diversity peaks at mid-latitudes and decreases monotonically towards low and high latitudes. The observed latitudinal diversity gradient is not unique to these invertebrates. Many macro- (e.g. Villagrán and Hinojosa 1997; Samaniego and Marquet 2009; Moreno et al. 2021) and microorganisms (i.e. protists; Fernández et al. 2016; Campello-Nunes et al. 2022) also exhibit a bell-shaped latitudinal diversity gradient in Chile. This finding suggests that millipede species richness covaries latitudinally with that of other Chilean taxa. Probably all Chilean taxa share a common biogeographic history determined, for example, by historical contingencies (desertification at low latitudes, glaciations at high latitudes) that originated Chile’s climatic gradient.

Millipedes are recognized for their low vagility and for occupying narrow geographic ranges (Sierwald and Bond 2007; Golovatch and Kime 2009). Consistently, we observed that most millipede species have narrow geographic ranges in Chile. Many of the species with narrow ranges accumulate in central Chile, contributing to the diversity peak we observed in mid-latitudes. This area is a known hotspot of diversity and endemism (Arroyo et al. 2004; Fernández 2010; Fernández et al. 2015; Campello-Nunes et al. 2022). Many of these organisms are threatened by habitat fragmentation and other human activities (Arroyo et al. 2004; Fernández et al. 2009) and possibly millipedes are no exception. The mid-latitudes harbor about 65 endemic millipede species with very narrow geographic ranges (Parra-Gómez submitted). Unfortunately, there is no information on the conservation status of Chilean millipedes. We presume that rare species, i.e. uncommon, scarce or infrequent species (e.g. *Siphonotus jacqueminae* Mauriès and Silva, 1971, *Myrmekia karykina* Attems, 1898, *Oligodesmus nitidus* Attems, 1898, *Polyxenus rossi* Chamberlin, 1957), are those facing the greatest risk of extinction in Chile.

Millipede diversity exhibited a positive correlation with a subset of climatic variables related to climatic seasonality, water availability and ambient energy inputs. Thus, millipedes are more diverse in latitudinal bands that exhibit a mesophilic or temperate climate (i.e. warm and humid) during most of the year, supporting two of the biogeographic hypotheses tested, i.e. climate stability and water-energy balance, respectively. In Chile, the mesophilic or temperate climate is present during most of the year in mid-latitudes (Veblen et al. 2007), explaining the diversity peak we have observed. Consistently, millipede diversity in other regions is also predicted by environmental variables used as surrogates for water-energy balance such as precipitation and temperature (Cooper 2022a; Cooper 2022b). Possibly the need for warm, humid climates is a phylogenetically conserved trait in millipedes (Kadamannaya et al. 2009; Cooper 2022b). Therefore, these arthropods may have evolutionary constraints that prevent them from adapting to new climates (Wiens et al. 2010; Fernández et al. 2022). Of course, this does not mean that there are no millipede species adapted to live in harsh climates. In Chile, the arid and polar climates of low and high latitudes, respectively, also harbor millipede species. However, the habitats of these areas are of post-Pleistocene origin and are therefore more recent than the habitats of mid-latitudes (Villagrán and Hinojosa 1997). These areas also have fewer millipede species than mid-latitudes. Therefore, the species composition present in high and low latitudes possibly represents cases of post-Pleistocene recolonization and/or cases of recent adaptation to severe climates. Among the adaptations that millipedes inhabiting the low and high latitudes of Chile may exhibit are a fossorial lifestyle and conglobation, which allow them to conserve moisture and survive in suboptimal sites (Golovatch 2009).

Beta diversity or spatial variation in millipede species composition varies from north to south based on strong species turnover compared to nestedness. Species turnover or species replacement is particularly strong and significant between the arid climate of low latitudes, the temperate climate of mid-latitudes, and the polar climate of high latitudes. Species turnover is often more important than nestedness when species distributions occur in response to an environmental gradient or when there are spatial and historical constraints (Baselga 2012; Fernández et al. 2016). Therefore, the observed beta diversity pattern supports the idea that millipede diversity is distributed from north to south according to the latitudinal gradient of climatic stability and water-energy balance. Likewise, the pattern of beta diversity supports the existence of evolutionary constraints (niche conservatism) that limit the adaptation of millipedes to the more severe climates of northern and southern Chile. Possibly, diversity is lower in low and high latitudes because historical contingencies that established severe climatic conditions in those areas limited the colonization of most mesophilic millipede species. At least this is the explanation that has been postulated to explain the low diversity exhibited by plants, animals, and protists in northern and southern Chile (Villagrán and Hinojosa 1997 Samaniego and Marquet 2009; Fernández 2015; Fernández et al. 2016; Campello-Nunes et al. 2022).

## 5. Conclusions

There are 96 native millipede species in Chile (considering described species, species that have not yet been formally described and subspecies that will be promoted to species), although we estimate that the diversity is between 125 and 197 species. On the other hand, our estimate is based on limited data so that the real diversity of Chilean millipedes could be much higher. In line with this conclusion, rarefaction and extrapolation curves suggest that the sampling effort invested in the last 175 years has been insufficient to record a substantial number of millipede species inhabiting Chile. To reverse this situation we need to significantly increase sampling effort across the extensive latitudinal and altitudinal gradients that characterize Chile.

The species richness of Chilean millipedes is distributed from north to south following a bell-shaped latitudinal diversity gradient, i.e. diversity peaks at mid-latitudes and decreases towards low and high latitudes. The diversity peak is caused by the accumulation of species with narrow geographic ranges in the mid-latitudes, a zone recognized as a hotspot of biodiversity and endemism for multi- and unicellular organisms. Species composition changes significantly between the arid climate of low latitudes, the temperate climate of mid-latitudes and the polar climate of high latitudes. Consistently, the variables that best predict the pattern and causes underlying the latitudinal diversity gradient of Chilean millipedes are climate stability, water availability and ambient energy inputs. Thus, Chilean millipedes are more diverse at sites that exhibit temperate (humid and warm) climates throughout much of the year. This result suggests that the biogeography of Chilean millipedes is predicted by the mechanisms proposed by two biogeographic hypotheses, i.e., climatic stability and water-energy balance. Possibly, water availability, ambient energy inputs, and climatic stability also predict broad-scale diversity patterns in millipedes from other regions of the planet.

## 7. Authors’ contributions

APG and LDF conceived the idea. APG collected the data and LDF conducted the statistical and biogeographical analyses. Both authors wrote the manuscript and gave final approval for publication.

## 8. Acknowledgments

APG thanks his parents for supporting his undergraduate studies in biology. LDF is funded by the Agencia Nacional de Investigación y Desarrollo (ANID), Chile, via FONDECYT projects 11170927 and 1220605. Both authors thank Jorge Pérez, Francisco Urra, Juan Campodonico and Edgardo Flores for providing biological material for this study.

## References

Adis J (2002) Amazonian Arachnida and Myriapoda identification keys to all classes, orders, families, some genera, and lists of know terrestrial species. Pensoft Publishers, Sofia, Bulgaria, 590 pp.

Arroyo MTK, Marquet PA, Marticorena C, Simonetti JA, Cavieres L, Squeo F, Rozzi R (2004) Chilean winter Rainfall–Valdivian forests. In: Mittermeier RA, Gil PR, Hoffmann M, Pilgrim J, Brooks T, Mittermeier CG, Lamoreux J, da Fonseca GAB (Eds), Hotspots revisted: Earth’s biologically wealthiest and most threatened ecosystems. CEMEX, México D.F, 99–103.

Attems CG (1898) System der Polydesmiden. I. Theil. Denkschriften der Kaiserlichen Akademie der Wissenschaften / Mathematisch-Naturwissenschaftliche Classe 67: 221–482.

Attems CG (1903) Beiträge zur myriopodenkunde. Zoologische Jahrbücher Abteilung für Systematik, Geographie und Biologie der Tiere 18: 63–154.

Baselga A (2012) The relationship between species replacement, dissimilarity derived from nestedness, and nestedness. Global Ecology and Biogeography 21(12): 1223–1232. https://doi.org/10.1111/j.1466-8238.2011.00756.x

Baselga A, Orme CDL (2012) betapart: an R package for the study of beta diversity. Methods in Ecology and Evolution 3(5): 808–812. https://doi.org/10.1111/j.2041-210X.2012.00224.x

Beck HE, Zimmermann NE, McVivar TR, Vergopolan N, Berg A, Wood EF (2018) Present and future Köppen-Geiger climate classification maps at 1-km resolution. Scientific Data 5(180214): 1–12. https://doi.org/10.1038/sdata.2018.214

Bivand R, Hauke J, Kossowski T (2013) Computing the Jacobian in Gaussian spatial autoregressive models: an illustrated comparison of available methods. Geographical Analysis 45(2): 150–179. https://doi.org/10.1111/gean.12008

Campello-Nunes PH, Woelfl S, da Silva-Neto ID, Paiva TS, Fernández LD (2022) Checklist, diversity and biogeography of ciliates (Ciliophora) from Chile. European Journal of Protistology 84(125892). https://doi.org/10.1016/j.ejop.2022.125892

Certes A (1889) Protozoaires. In: Mission Scientifique du Cap Horn 1882–1883. Zoologie. Paris, 1–53. Available from: https://doi.org/10.5962/bhl.title.2480.

Chamberlin RV (1957) The Diplopoda of the Lund University and California Academy of Sciences expeditions. Lunds Universitets Arsskrift, N. Följd, Avd. 2: 1–44.

Chao A, Jost L (2015) Estimating diversity and entropy profiles via discovery rates of new species. Methods in Ecology and Evolution 6(8): 873–882. https://doi.org/10.1111/2041-210X.12349

Chao A, Gotelli NJ, Hsieh TC, Sander EL, Ma KH, Colwell RK, Ellison AM (2014) Rarefaction and extrapolation with Hill numbers: a framework for sampling and estimation in species diversity studies. Ecological Monographs 84(1): 45–67. https://doi.org/10.1890/13-0133.1

Clarke KR, Ainsworth M (1993) A method of linking multivariate community structure to environmental variables. Marine Ecology Press Series 92: 205–219. https://doi.org/10.3354/meps092205

Clarke KR, Gorley RN, Somerfield PJ, Warwick RM (2005) Change in marine communities: an approach to statistical analysis and interpretation, 3rd edn. PRIMER–E Ltda., Plymouth, UK.

Clarke KR, Gorley RN (2006) PRIMER v.6: User Manual / Tutorial. PRIMER–E Ltda., Plymouth, UK.

Cooper M (2022a) Assessment of latitudinal gradient in species richness of Sphaerotherium. New Visions in Biological Science 9: 14–20. https://doi.org/10.9734/bpi/nvbs/v9/1885A

Cooper M (2022b) The inverse latitudinal gradient in species richness of forest millipedes: Centrobolus Cook, 1897. New Visions in Biological Science 9: 82–88. https://doi.org/10.9734/bpi/nvbs/v9/1898A

Currie DJ (1991) Energy and large-scale patterns of animal- and plant-species richness. The American Naturalist 137(1): 27–49.

Davies BJ, Darvill CM, Lovell H, Bendle JM, Dowdeswell JA, Fabel D, García JL, Geiger A, Glasser NF, Gheorghiu DM, Harrison S, Hein AS, Kaplan MR, Martin JRV, Mendelova M, Palmer A, Pelto M, Rodés Á, Sagredo EA, Smedley RK, Smellie JL, Thorndycraft VR (2020) The evolution of the Patagonian Ice Sheet from 35 ka to the present day (PATICE). Earth-Science Reviews 204(103152): 1–77. https://doi.org/10.1016/j.earscirev.2020.103152

Demange J-M, Silva F (1971a) Nouvelle espèce chilienne du genre Autostreptus Silvestri et description du matériel type de Iulus chilensis Gervais, 1847, type du genre (Myriapode, Diplopode, Spirostreptoidea, Spirostreptidae, Spirostreptinae). Bulletin du Muséum National d’Histoire Naturelle, 2e série 42(4): 708–715.

Demange J-M, Silva F (1971b) Abatodesmus velosoi nov. sp., nouvelle espèce chilienne de la famille des Sphaerotrichopidae (Myriapode, Diplopodae: Polydesmoidea). Bulletin du Muséum National d’Histoire Naturelle 42(5): 881–886.

Demange J-M, Silva F (1976a) Etudes de quelques genres chiliens de la famille des Sphaerotrichopidae (Myriapodes, Diplopodes, Polydesmoidae). Bollettino del Laboratorio di Entomologia Agraria “Filippo Silvestri” di Portici 33: 15–33.

Demange J-M, Silva F (1976b) Contribution á la connaissance des espèces originaires du Chili par F. Silvestri en 1902. Bollettino del Laboratorio di Entomologia Agraria “Filippo Silvestri” di Portici 33: 34–43.

Evans KL, Warren PH, Gaston KJ (2005) Species-energy relationships at the macroecological scale: a review of the mechanisms. Biological Reviews 80(1): 1–25. https://doi.org/10.1017/S1464793104006517

Fernández L (2010) Foraminíferos (Protozoa: Foraminiferida) del estuario del río Contaco (40° 33’S; 73° 43’O), Chile. Boletín de Biodiversidad de Chile 4: 18–62.

Fernández L (2011) Nuevos registros y ampliaciones de rango ¿Para qué? Boletín de Biodiversidad de Chile 5: 1–2.

Fernández LD (2015) Source-sink dynamics shapes the spatial distribution of soil protists in an arid shrubland of northern Chile. Journal of Arid Environments 113: 121–125. https://doi.org/10.1016/j.jaridenv.2014.10.007

Fernández L, Rau J, Arriagada A (2009) Calidad de la vegetación ribereña del río Maullín (41 28’S; 72 59’O) utilizando el índice QBR. Gayana Botanica 66(2): 269–278. https://doi.org/10.4067/S0717-66432009000200011

Fernández LD, Lara E, Mitchell EAD (2015) Checklist, diversity and distribution of testate amoebae in Chile. European Journal of Protistology 51(5): 409–424. https://doi.org/10.1016/j.ejop.2015.07.001

Fernández LD, Fournier B, Rivera R, Lara E, Mitchell EAD, Hernández CE (2016) Water–energy balance, past ecological perturbations and evolutionary constraints shape the latitudinal diversity gradient of soil testate amoebae in south-western South America. Global Ecology and Biogeography 25(10): 1216–1227. https://doi.org/10.1111/geb.12478

Fernández LD, Seppey CVW, Singer D, Fournier B, Tatti D, Mitchell EAD, Lara E (2022) Niche conservatism drives the elevational diversity gradient in major groups of free-living soil unicellular eukaryotes. Microbial Ecology 83(2): 459–469. https://doi.org/10.1007/s00248-021-01771-2

Gervais P (1847) Myriapodes. In: Walckenaer CA, Gervais P (Eds), Histoire naturelle des Insectes. Aptères. Librairie Encyclopédique de Roret, Paris, 1–330. Available from: https://doi.org/10.5962/bhl.title.61095.

Golovatch SI (2014) On some new or poorly-known millipedes from Chile and Argentina (Diplopoda). Russian Entomological Journal 23: 249–281. https://doi.org/10.15298/RUSENTJ.23.4.02

Golovatch SI (2009) Millipedes (Diplopoda) in extreme environments. In: Golovatch SI, Makarova OL, Babenko AB, Penev LD (Eds), Species and Communities in Extreme Environments: Festschrift towards the 75th Anniversary and a Laudatio in Honour of Academican Yuri Ivanovich Chernov. Pensoft & KMK Scientific Press, Sofia-Moscow, 87–112.

Golovatch SI, Kime RD (2009) Millipede (Diplopoda) distributions: a review. Soil Organisms 81(3): 565–597.

Gregory-Wodzicki KM (2000) Uplift history of the Central and Northern Andes: A review. Geological Society of America Bulletin 112(7): 1091–1105. https://doi.org/10.1130/0016-7606(2000)112<1091:UHOTCA>2.0.CO;2

Hammer Ø, Harper DAT, Ryan PD (2001) PAST: paleontological statistics software for education and data analysis. Paleontologia Electronica 4(1): 1–9.

Hawkins BA, Field R, Cornell HV, Currie DJ, Guégan J-F, Kaufman DM, Kerr JT, Mittelbach GG, Oberdorff T, O’Brien EM, Porter EE, Turner JRG (2003) Energy, water, and broad-scale geographic patterns of species richness. Ecology 84(12): 3105–3117. https://doi.org/10.1890/03-8006

Hsieh TC, Ma KH, Chao A (2016 iNEXT: an R package for rarefaction and extrapolation of species diversity (Hill numbers). Methods in Ecology and Evolution 7(12): 1451–1456

Jung W (1942) Südchilenische Thekamöeben. Archiv für Hydrobiologie 95: 253–356.

Kadamannaya BS, Sridhar KR, Sahadevan S (2009) Seasonal periodicity of pill millipedes (Arthrosphaera) and earthworms of the Western Ghats, India. World Journal of Zoology 4(2): 63–69.

Kerr JT, Packer L (1997) Habitat heterogeneity as a determinant of mammal species richness in high-energy regions. Nature 385: 252–254. https://doi.org/10.1038/385252a0

Korsós Z, Read H (2012) Redescription of Zinagon chilensis (Silvestri, 1903) from Chile, with a species list of Iulomorphidae from the Southern Hemisphere (Diplopoda: Spirostreptida: Epinannolenidea). Zootaxa 3493: 38–48. https://doi.org/10.11646/zootaxa.3493.1.4

Krabbe E (1982) Systematik der Spirostreptidae (Diplopoda, Spirostreptomorpha). Abhandlungen und Verhandlungen des Naturwissenschaftlichen Vereins in Hamburg (N.F.) 24: 1–476.

Marek PE, Buzatto BA, Shear WA, Means JC, Black DG, Harvey MS, Rodriguez J (2021) The first true millipede—1306 legs long. Scientific Reports 11(23126): 8. https://doi.org/10.1038/s41598-021-02447-0

Mauriès J-P, Silva F (1971) Colobognathes du Chili. I. Espèces nouvelles du genre Siphonotus Brandt (Diplopoda). Bulletin du Muséum National d’Histoire Naturelle 2(42): 887–902.

McCain CM (2009) Global analysis of bird elevational diversity. Global Ecology and Biogeography 18(3): 346–360. https://doi.org/10.1111/j.1466-8238.2008.00443.x

Mesibov R (2017) New records for millipedes from southern Chile (Polydesmida: Dalodesmidae; Polyzoniida: Siphonotidae). Biodiversity Data Journal 5: 1–5. https://doi.org/10.3897/BDJ.5.e15919

Minelli A (2015) Treatise on Zoology - Anatomy, Taxonomy, Biology. The Myriapoda, Volume 2. Brill, Leiden, The Netherlands, 482 pp. Available from: https://doi.org/10.1163/9789004188273.

Moreno RA, Labra FA, Cotoras DD, Camus PA, Gutiérrez D, Aguirre L, Rozbaczylo N, Poulin E, Lagos NA, Zamorano D, Rivadeneira MM (2021) Evolutionary drivers of the hump-shaped latitudinal gradient of benthic polychaete species richness along the Southeastern Pacific coast. PeerJ 9(e12010): 1–24. https://doi.org/10.7717/peerj.12010

Parra-Gómez A (2022) Records about the alien millipede Oxidus gracilis (C. L. Koch, 1847) (Diplopoda: Polydesmida: Paradoxosomatidae) in continental Chile. Revista Chilena de Entomología 48(1): 73–79. https://doi.org/10.35249/rche.48.1.22.06

Parra-Gómez A, Faúndez EI (2021) First teratological case in Pleonaraius pachyskeles Attems, 1898 (Polydesmida: Dalodesmidae) including new distributional records in Chile. Boletín del Museo Nacional de Historia Natural del Paraguay 25(2): 144–149.

Pianka ER (1966) Latitudinal gradients in species diversity: a review of concepts. The American Naturalist 100(910): 33–46. https://doi.org/10.1086/282398

Rodríguez MÁ, Belmontes JA, Hawkins BA (2005) Energy, water and large-scale patterns of reptile and amphibian species richness in Europe. Acta Oecologica 28(1): 65–70. https://doi.org/10.1016/j.actao.2005.02.006

Samaniego H, Marquet PA (2009) Mammal and butterfly species richness in Chile: taxonomic covariation and history. Revista Chilena de Historia Natural 82: 135–151.

Shear WA (1988) The Chordeumatid millipeds of Chile (Diplopoda, Chordeumatida). American Museum Novitates (2912): 1–10. https://doi.org/2246/5171

Shelley RM, Golovatch SI (2011) Atlas of Myriapod Biogeography. I. Indigenous ordinal and supra-ordinal distributions in the Diplopoda: perspectives on taxon origins and ages, and a hypothesis on the origin and early evolution of the class. Insecta Mundi 0158: 1–134.

Short M, Vahtera V (2017) Phylogenetic relationships of millipedes in the subclass Penicillata (Diplopoda) with a key to the genera. Journal of Natural History 51(41–42): 2443–2461. https://doi.org/10.1080/00222933.2017.1380241

Sierwald P, Bond JE (2007) Current status of the myriapod class Diplopoda (Millipedes): taxonomic diversity and phylogeny. Annual Review of Entomology 52(1): 401–420. https://doi.org/10.1146/annurev.ento.52.111805.090210

Silva F, Vivar C (1973) Presencia de Chilorus zapallar Chamberlin, 1957, en el bosque nativo de Quintero. Anales Del Museo de Historia Natural de Valparaíso 6: 217–223.

Silva F, Vivar C (1974) Miriápodos: I diplópodos del Parque Nacional “Vicente Perez Rosales.” Anales del Museo de Historia Natural de Valparaíso 7: 285–291.

Silva F, Sáiz F (1975) Investigaciones ecológicas de los diplópodos del Parque Nacional “Fray Jorge.” Anales del Museo de Historia Natural de Valparaíso 8: 17–28.

Silva F, Veloso A, Solervicens J, Ortiz JC (1968) Investigaciones zoológicas en el Parque Nacional Vicente Pérez Rosales y zona de Pargua. Noticiario Mensual del Museo Nacional de Historia Natural: 3–12.

Spelda J (2015) The southernmost millipedes found on Guarello Island, southern Chile (Diplopoda, Polydesmida, Dalodesmidae). Spixiana 38(2): 186.

Urzua A, Silva F (1981) Componentes químicos de las secreciones defensivas de miriápodos-diplópodos del género Autostreptus, Silvestri, 1905 (Spirostreptida-Spirostreptidae). Anales del Museo de Historia Natural de Valparaíso 14: 271–273.

Veblen TT, Young KR, Ome AR (2007) The physical geography of South America. Oxford University Press, Oxford, 361 pp. Available from: 10.1093/oso/9780195313413.001.0001.

Vega-Román E, Ruiz V, Arancibia-Ávila P (2019) First record of the genus Polyxenus Latreille (Diplopoda: Penicillata: Polyxenida) in the supralittoral zone of Cocholgüe, Biobío Region, Chile. Revista Chilena de Entomología 45(3): 399–402. https://doi.org/10.35249/rche.45.3.19.11

Villagrán C, Hinojosa LF (1997) Historia de los bosques del sur de Sudamérica, II: Análisis fitogeográfico. Revista Chilena de Historia Natural 70: 241–267.

Wiens JJ, Ackerly DD, Allen AP, Anacker BL, Buckley LB, Cornell HV, Damschen EI, Jonathan Davies T, Grytnes J-A, Harrison SP, Hawkins BA, Holt RD, McCain CM, Stephens PR (2010) Niche conservatism as an emerging principle in ecology and conservation biology. Ecology Letters 13(10): 1310–1324. https://doi.org/10.1111/j.1461-0248.2010.01515.x

